# Minus the Error: Testing for Positive Selection in the Presence of Residual Alignment Errors

**DOI:** 10.1101/2024.11.13.620707

**Authors:** Avery Selberg, Nathan L Clark, Timothy B Sackton, Spencer V Muse, Alexander G Lucaci, Steven Weaver, Anton Nekrutenko, Maria Chikina, Sergei L. Kosakovsky Pond

## Abstract

Positive selection is an evolutionary process which increases the frequency of advantageous mutations because they confer a fitness benefit. Inferring the past action of positive selection on protein-coding sequences is fundamental for deciphering phenotypic diversity and the emergence of novel traits. With the advent of genome-wide comparative genomic datasets, researchers can analyze selection not only at the level of individual genes but also globally, delivering systems-level insights into evolutionary dynamics. However, genome-scale datasets are generated with automated pipelines and imperfect curation that does not eliminate all sequencing, annotation, and alignment errors. Positive selection inference methods are highly sensitive to such errors. We present BUSTED-E: a method designed to detect positive selection for amino acid diversification while concurrently identifying some alignment errors. This method builds on the flexible branch-site random effects model (BUSTED) for fitting distributions of dN/dS, with a critical modification: it incorporates an “error-sink” component to represent an abiological evolutionary regime. Using several genome-scale biological datasets that were extensively filtered using state-of-the art automated alignment tools, we show that BUSTED-E identifies pervasive residual alignment errors, produces more realistic estimates of positive selection, reduces bias, and improves biological interpretation. The BUSTED-E model promises to be a more stringent filter to identify positive selection in genome-wide contexts, thus enabling further characterization and validation of the most biologically relevant cases.

## Introduction

Comparative analyses that estimate evolutionary parameters from multiple sequence alignments (MSAs) are a cornerstone of many genomic investigations. For example, conservation metrics, such as those produced by the phyloP software, are widely used to pinpoint functionally constrained regions, guide variant interpretation, and identify elements under purifying selection (***Pollard et al., 2010***). A counterpart of purifying selection is episodic diversifying selection (EDS). Whereas purifying selection eliminates deleterious mutations, maintaining sequence similarity across large evolutionary distances, EDS is the process in which advantageous genetic changes are repeatedly fixed along phylogenetic branches, contributing to between-species sequence divergence. Typically, only a small fraction of codons (or amino acid residues) in a gene are subject to selection during these episodes. The ability to identify EDS with reasonable specificity is a necessary condition for more complex analyses which seek to connect adaptive changes to emergence of novel phenotypes or other evolutionary processes.

Modern comparative genomics covers hundreds of species and tens of thousands of genes. These large-scale datasets typically rely on automated pipelines to assemble and align sequences. Despite continuing improvements in alignment algorithms, it remains nearly impossible to guarantee error-free alignments, especially when manual curation is impractical. These alignment errors present a significant obstacle for EDS analyses, which rely on distinguishing synonymous and nonsynonymous coding substitutions (i.e., dN/dS analyses). Even low rates of alignment error can profoundly bias inference of EDS (***Schneider et al., 2009***). Further, while estimates of purifying selection improve as more species are added (***zoo, 2020***), peformance of dN/dS tests for EDS can actually degrade as additional sequences increase the probability of a local alignment error. This can undermine biological interpretations.

Previous studies using simulated alignment errors have demonstrated increased false positive rates and assessed mitigation strategies (***Fletcher and Yang, 2010; Privman et al., 2012; Jordan and Goldman, 2012***). One consistent conclusion in simulated alignment error studies is that PRANK-C codon alignments were the least affected and could not be further improved by standard filtering programs (***Jordan and Goldman, 2012***). Redelings et al. (***Redelings, 2014***) proposed a method for jointly estimating both the positive selection signal and alignment errors in a Bayesian framework. This approach improved alignment accuracy, even for PRANK-C alignments, achieving performance close to that of error-free data. However, the computational cost of Markov Chain Monte Carlo sampling in this framework renders it impractical for genome-scale analyses, and this approach has seen limited adoption.

Here, we introduce BUSTED-E, a method designed to overcome some of the challenges of quantifying EDS in the presence of alignment errors at a minimal additional computational cost. Based on the BUSTED framework, BUSTED-E incorporates an “error-sink” component that captures aberrant evolutionary patterns unrelated to true biological processes. This novel approach flags problematic alignment regions, refines estimates of selection, and reduces spurious signals commonly encountered in large-scale comparative analyses. By enhancing the robustness of alignment-dependent inferences, BUSTED-E yields more reliable insights into the evolutionary forces shaping genomic diversity.

## Results

To develop intuition for how BUSTED-E works, consider deciding if a gene experienced EDS using a method that estimates dN/dS (*ω*) and uses a statistical test to obtain a p-value for the hypothesis that *ω >* 1 on some sites in the MSA and/or branches of the corresponding phylogenetic tree.

The new BUSTED-E method is based on BUSTED (***Wisotsky et al., 2020***), including synonymous rate variation, which estimates the *ω* distribution across sites with three classes (*ω*_1_, *ω*_2_, and *ω*_3_; the number of classes is a user-tunable parameter). Evolution along each branch at each site is parametrized by a random independent draw from this distribution, and only one value (*ω*_3_) is permitted to take values g 1, i.e., positive diversifying selection for amino acid replacement.

BUSTED uses a likelihood ratio test (LRT) to compute a p-value versus the null hypothesis where *ω*_3_ f 1. Other frequently used models include those in the PAML suite (***Yang, 2007***) or Bayesian approaches (***Murrell et al., 2013; Rodrigue and Lartillot, 2014; Redelings, 2014***). The advantage of the BUSTED framework is that its modular organization can accommodate flexible rate distributions, while being computationally tractable and scalable (***Kosakovsky Pond et al., 2011***). BUSTED-E adds a category described by *ω*_*E*_ g 100, whose maximum weight is 1%. This “error-sink” category is meant to capture false signals derived from local alignment errors like those shown in ***Figure 1***. The bounds on the *ω*_*E*_ category are heuristic, adjustable, and informed by parameter estimates from large collections of alignments.

**Figure 1.**
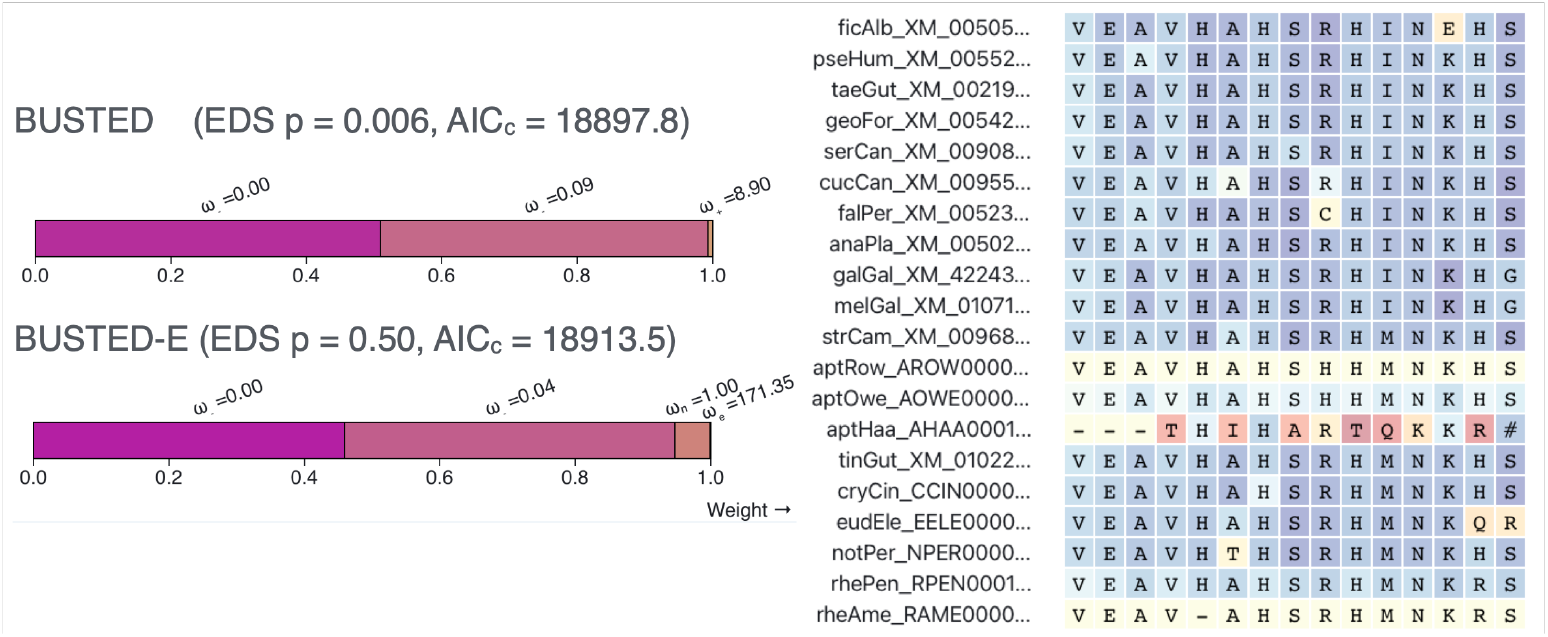
BUSTED and BUSTED-E applied to the *UROD* gene alignment of 40 bird sequences from Shultz and Sackton ***Shultz and Sackton (2019)*** which used a sophisticated alignment curation and masking procedure to remove artifacts. BUSTED infers three *ω* classes, including a class with *ω >* 1, and deduces the presence of episodic diversifying selection (EDS), *p* = 0.006. BUSTED-E introduces an additional error class (*ω >* 100, maximum weight f 1%). This changes improves model fit (small sample AIC), places a 0.1% fraction of the alignment in the error class, removes the distribution class with 1 *< ω <* 100, and eliminates evidence for EDS (*p* = 0.50). The error attribution procedure of BUSTED-E highlights (red colors) one problematic part of this alignment, located at the 3’ end of the gene, in the *Apteryx rowi (aptHaa)* sequence. BUSTED misinterprets such local misalignments as evidence of rapid diversification, whereas BUSTED-E has the option to correctly treat them as “noise”. Blue and yellow backgrounds represent low error probability. **Figure 1—figure supplement 1**. Apparent local misalignment at the 3’ end of the PXDNL gene from Shultz and Sackton (***Shultz and Sackton, 2019***). The color palette indicates empirical Bayes factor support for assignment to the error class.

To illustrate the new model we analyzed the uroporphyrinogen decarboxylase (UROD) gene involved in heme production. UROD is one of 11,267 genes analyzed for selection by Shultz and Sackton (***Shultz and Sackton, 2019***). Seeking to quantify positive selection across bird and mammalian genomes, their work used a sophisticated pipeline that included multiple filtering steps for the MSAs and multiple positive selection inference methods. Notably, the MSAs were produced using PRANK-C, widely assumed to produce the most accurate alignments for positive selection inference (***Fletcher and Yang, 2010; Markova-Raina and Petrov, 2011; Jordan and Goldman, 2012***). BUSTED identifies significant evidence of EDS (*p* = 0.006) in UROD (***Figure 1***) with a moderate positive selection component *ω*_3_ = 8.90 with localized support (0.7% of the alignment). Manual inspection of the MSA reveals local misalignment near the 3’ end, resulting in several apparent multi-nucleotide substitutions and codon sites with low homology (***Figure 1***). BUSTED-E estimates the same three *ω* classes, with the addition of the error class *ω*_*E*_, which is forced to have *ω* g 100 and cannot draw more than 1% weight. This leads to a dramatic reversal in support for EDS for UROD (*p* = 0.50), where all the apparent positive selection weight is absorbed by the error class, and *ω*_3_ = 1(5.2%) now reflects neutral evolution. As a point of reference, a previous comparative study of UROD and other heme production genes found them to be evolving under strong purifying selection in a broader taxonomic context (***Tzou et al., 2014***).

### Reanalysis of genome-wide scans

We selected four studies (***Schneider et al., 2009; Nguyen et al., 2015; Wu et al., 2018; Shultz and Sackton, 2019***) with publicly available alignments (***Table 1***, Supplementary Table 1). All four used one or more dN/dS tests (implemented in HyPhy and/or PAML) to examine a thousand or more coding sequence alignments for evidence of EDS. Each study employed automated quality control and filtering, reflecting a field-wide appreciation that such filtering is necessary. The studies spanned a decade, were done by unconnected research groups, varied in taxonomic range and alignment size, and differed in research goals. All original and filtered multiple sequence alignments and trees are available from https://github.com/veg/pub-data.

**Table 1.**
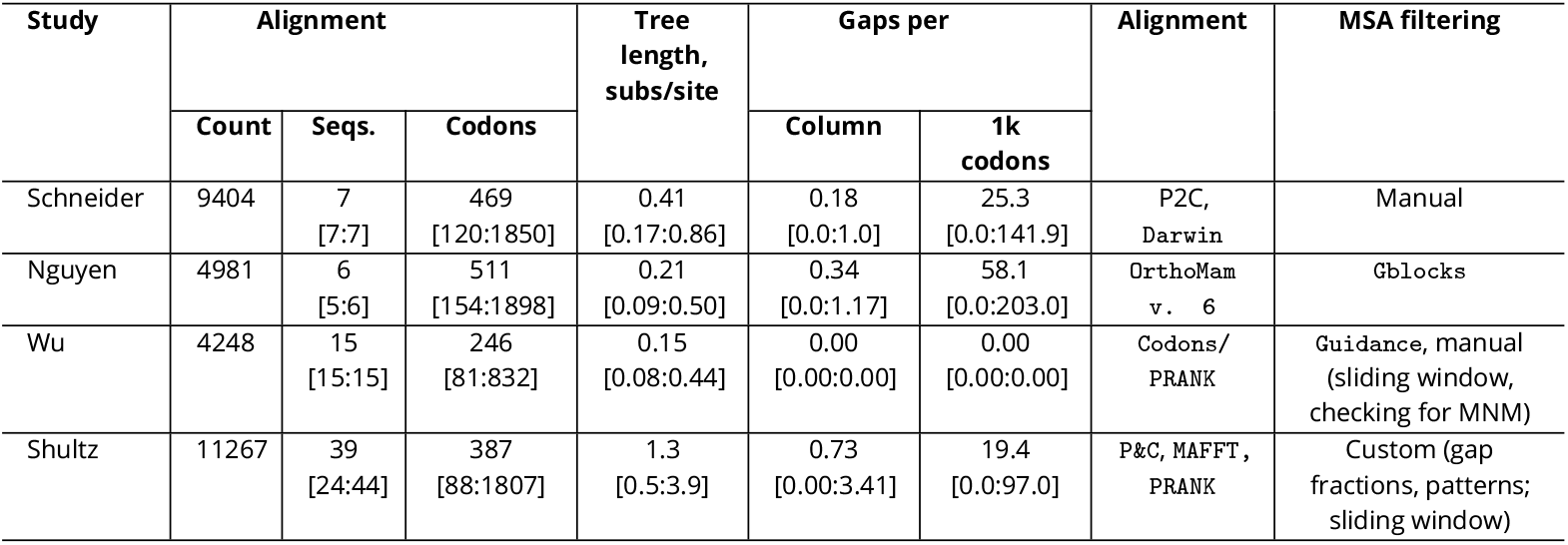
The large scale datasets used for BUSTED-E evaluation. For all per-alignment quantities (sequences, codons, tree length, gaps per) we report the median value and the [2.5% : 97.5%] range. Gaps per sequence are normalized per 1000 codons. Tree length is the cumulative branch length (expected substitutions/nucleotide) under the simple (MG94xREV) codon model. Abbreviations for the Alignment column are as follows. P2C: translated protein sequences are aligned, then mapped back to codon sequences; P&C: protein based homology and filtering, codon-level alignment; MNM : multi-nucleotide mutations.

### BUSTED-E drastically reduces the rate of likely erroneous positive selection inference

Compared to BUSTED (***Wisotsky et al., 2020***) with synonymous rate variation, error-compensated BUSTED-E model returned many fewer positive EDS results (***Table 2***) across the range of test stringencies (***Figure 2***). Even when using the less conservative model averaged approach (MA), where results from BUSTED-S and BUSTED-E models are combined to mitigate the potential loss of power (see Methods), the EDS detection rate is substantially lower. Note that if only a very small fraction of alignments are truly subject to positive selection, then the detection rate should be close to the chosen significance level, assuming the method is neither too conservative nor too anti-conservative. When using a significance level of 0.01 we expect approximately 1% of alignments to be identified as positively selected, since 0.01 is the expected false positive rate.

**Table 2.**
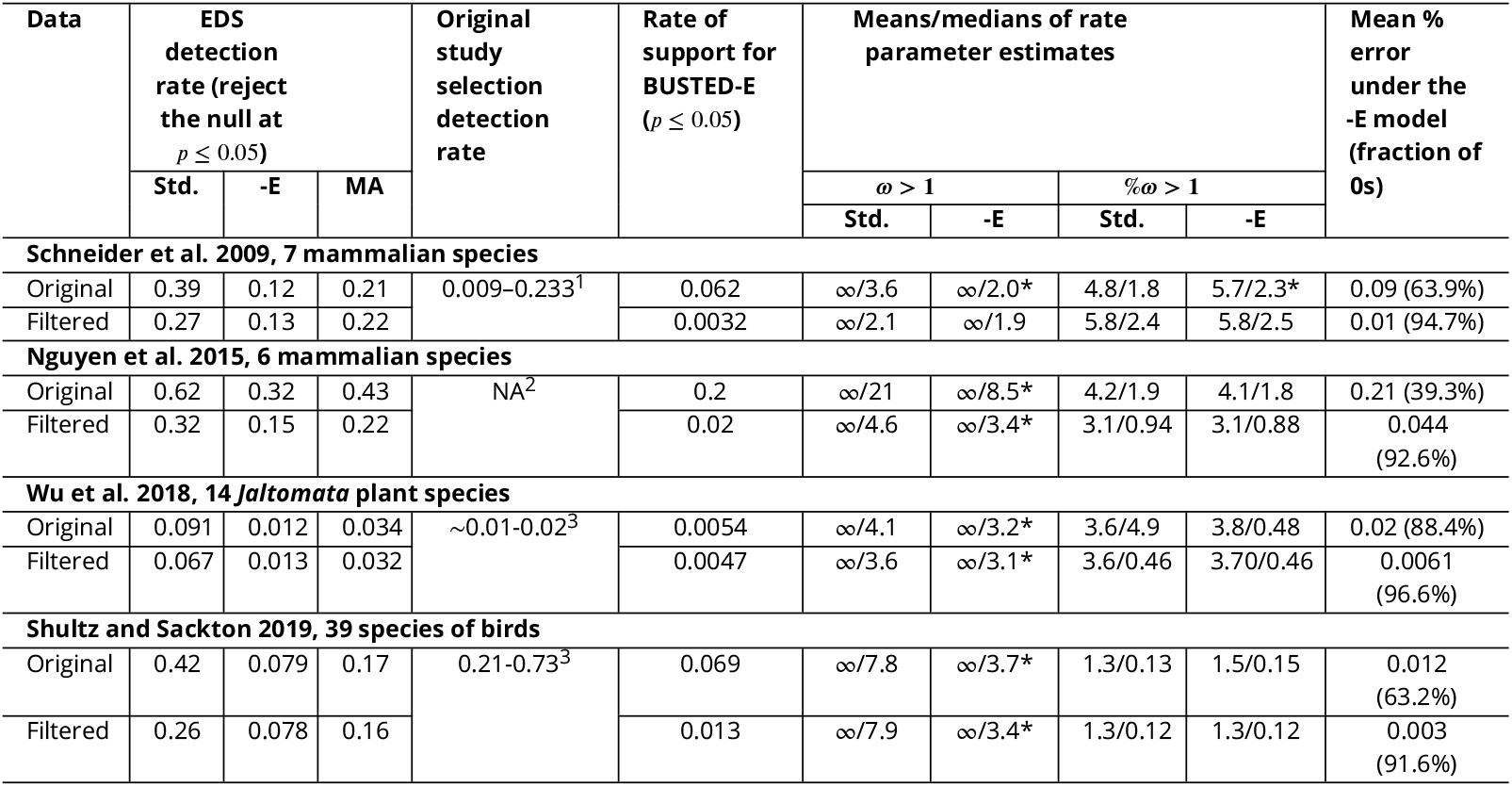
Comparative Method Performance on Genome-Scale Datasets. BUSTED (Std.) and BUSTED-E (-E) performance on empirical data sets; both original and filtered using BUSTED-E (see text). MA: model average. *: estimates between BUSTED-E and BUSTED have a Mann-Whitney test p *<* 0.0001. (1) Per lineage branch-site tests using PAML at FDRf0.05 following the Rom correction. (2) No direct dN/dS ë 1 tests were done. (3) FDRf0.01 corrected clade branch-site tests using PAML for different clades. (4). p f 0.05 for different PAML model pairs and the original BUSTED method. The study used a 0.14 rate by requiring unanimous method consent.

| Data | EDS<br>detection<br>rate (reject<br>the null at<br>$p \leq 0.05$) | | | Original<br>study<br>selection<br>detection<br>rate | Rate<br>of<br>support<br>for<br>BUSTED-E<br>($p \leq 0.05$) | Means/medians of rate<br>parameter estimates | | | | Mean %<br>error<br>under the<br>-E model<br>(fraction of<br>0s) |
| --- | --- | --- | --- | --- | --- | --- | --- | --- | --- | --- |
| | Std. | -E | MA | | | $\omega > 1$ | | %$\omega > 1$ | | |
|  |  |  |  |  |  | Std. | -E | Std. | -E |  |
| Schneider et al. 2009, 7 mammalian species |  |  |  |  |  |  |  |  |  |  |
| Original | 0.39 | 0.12 | 0.21 | 0.009–0.233<sup>1</sup> | 0.062 | $\infty/3.6$ | $\infty/2.0^*$ | 4.8/1.8 | 5.7/2.3* | 0.09 (63.9%) |
| Filtered | 0.27 | 0.13 | 0.22 | | 0.0032 | $\infty/2.1$ | $\infty/1.9$ | 5.8/2.4 | 5.8/2.5 | 0.01 (94.7%) |
| Nguyen et al. 2015, 6 mammalian species |  |  |  |  |  |  |  |  |  |  |
| Original | 0.62 | 0.32 | 0.43 | NA<sup>2</sup> | 0.2 | $\infty/21$ | $\infty/8.5^*$ | 4.2/1.9 | 4.1/1.8 | 0.21 (39.3%) |
| Filtered | 0.32 | 0.15 | 0.22 | | 0.02 | $\infty/4.6$ | $\infty/3.4^*$ | 3.1/0.94 | 3.1/0.88 | 0.044<br>(92.6%) |
| Wu et al. 2018, 14 *Jaltomata* plant species |  |  |  |  |  |  |  |  |  |  |
| Original | 0.091 | 0.012 | 0.034 | $\sim 0.01$–0.02<sup>3</sup> | 0.0054 | $\infty/4.1$ | $\infty/3.2^*$ | 3.6/4.9 | 3.8/0.48 | 0.02 (88.4%) |
| Filtered | 0.067 | 0.013 | 0.032 | | 0.0047 | $\infty/3.6$ | $\infty/3.1^*$ | 3.6/0.46 | 3.70/0.46 | 0.0061<br>(96.6%) |
| Shultz and Sackton 2019, 39 species of birds |  |  |  |  |  |  |  |  |  |  |
| Original | 0.42 | 0.079 | 0.17 | 0.21–0.73<sup>3</sup> | 0.069 | $\infty/7.8$ | $\infty/3.7^*$ | 1.3/0.13 | 1.5/0.15 | 0.012<br>(63.2%) |
| Filtered | 0.26 | 0.078 | 0.16 | | 0.013 | $\infty/7.9$ | $\infty/3.4^*$ | 1.3/0.12 | 1.3/0.12 | 0.003<br>(91.6%) |

**Figure 2.**
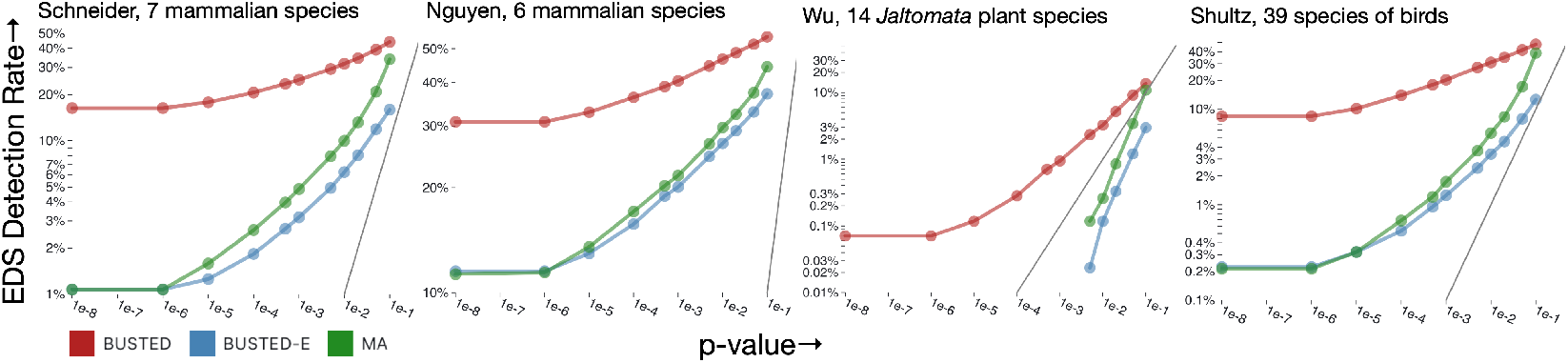
The rate at which genes are found to be subject to EDS as function of the nominal p-value cutoff. The grey line shows the expectation under the strict null (no positive selection, at most neutral evolution).

In ***Figure 2***, we observe that the significance rates for error-corrected models align more closely with expected patterns. Notably, three out of the four datasets exhibit a substantial proportion (∼10-30%) of alignments where BUSTED detects EDS, even for the smallest p-value thresholds. This suggests that such “certain” calls of EDS may stem from the presence of anomalous regions within many of these alignments that are assigned to the error class by BUSTED-E.

### BUSTED-E reduces bias in positive selection calls, improving biological interpretation

Because ground truth in selection analyses is unknown, we cannot claim that our method is provably better for detection of EDS. However, we can evaluate how the choice of method for genomewide analysis of positive selection impacts signal distribution.

Many genome-wide analyses aim to identify the functional contexts experiencing positive selection. To see how BUSTED-E might impact such inference, we ran enrichment analyses using a collection of functional annotation sources (KEGG, Reactome, GO, WikiPathways) on the top 5% (ranked by LRT) of genes (***Figure 3***). Analyzing a combined set of pathways with FDR *<* 0.2 for encrichment in either set, we find that BUSTED-E and BUSTED highlight different pathways with slightly more pathways (10 vs 8) being more strongly enriched in BUSTED-E. However, the results obtained by BUSTED-E are much more closely aligned with both the original Schulz and Sackton analyses, which made a conservative consensus call of several methods, and other analyses, highlighting the general overrepresentation of immune genes in the positively selected set (***Vinkler et al., 2023***). These findings are also consistent with the central role of host-pathogen interactions in driving EDS (***Enard et al., 2016***). Moreover, pathways enriched in BUSTED have little biological coherence or alignment with prior knowledge, which would be consistent with error contamination. Further analysis suggests factors other than the action of positive selection may contribute to EDS detection with BUSTED. Specifically, we observe that for pathways enriched in BUSTED the genes driving enrichment tend to be unusually long. To quantify the overall trend, we color code each pathway according to how well protein length alone predicts whether a gene is part of the foreground set identified by the method showing the highest enrichment for that pathway (***Figure 3***). We quantify the association via a rank-based AUC metric, where a value of 1 would indicate that the pathway genes are ranked entirely at the top by length while a value of 0.5 indicates a random length distribution. This approach reveals that for pathways specific to BUSTED, the genes contributing most to the enrichment exhibit length bias, i.e., longer proteins may disproportionately drive enrichment signals. For example, considering the top BUSTED pathway of “Microtubule-Based Transport” (GO:0099111) the median length for the foreground gene is 1593 codons while the overall median length in the dataset is 390. In contrast, the pathways highlighted by BUSTED-E have relatively weaker length bias with lower length AUCs. Some length bias is expected even under error-free conditions as greater alignment length increases sample size and power.

**Figure 3.**
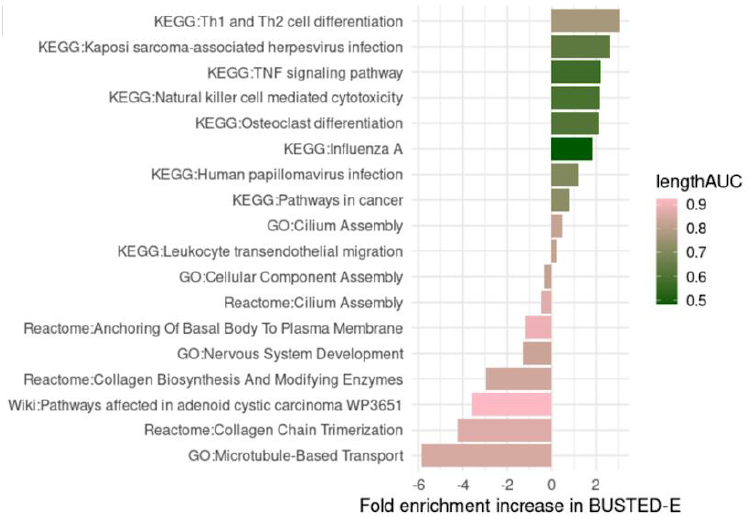
Enrichment/depletion analysis using top 5% LRT genes (BUSTED-E) as the foreground set, at FDR f 0.2. Each bar is colored by the AUC of a simple predictive model which uses alignment length to predict BUSTED LRT model ranking (see text).

### Datasets where BUSTED and BUSTED-E disagree have a characteristic pattern of inferred rates

***Figure 4*** shows a characteristic “shift” in both *ω*_3_ and its proportion among the genes where BUSTED and BUSTED-E disagree. BUSTED-E infers smaller *ω*_3_ values and larger corresponding weights when the two methods disagree (with BUSTED-E finding no evidence of selection in nearly all such cases). On the other hand, the distributions of *ω*_3_ and its weight are largely congruent for the genes where the two methods agree. While there are several distinct reasons why genes may be discordantly classified (see below), the major driver is the absorption of a small fraction of the alignment where very large values of *ω* find support into the error class, and the corresponding reduction in the *ω* estimates. For a concrete example, see ***Figure 1***.

**Figure 4.**
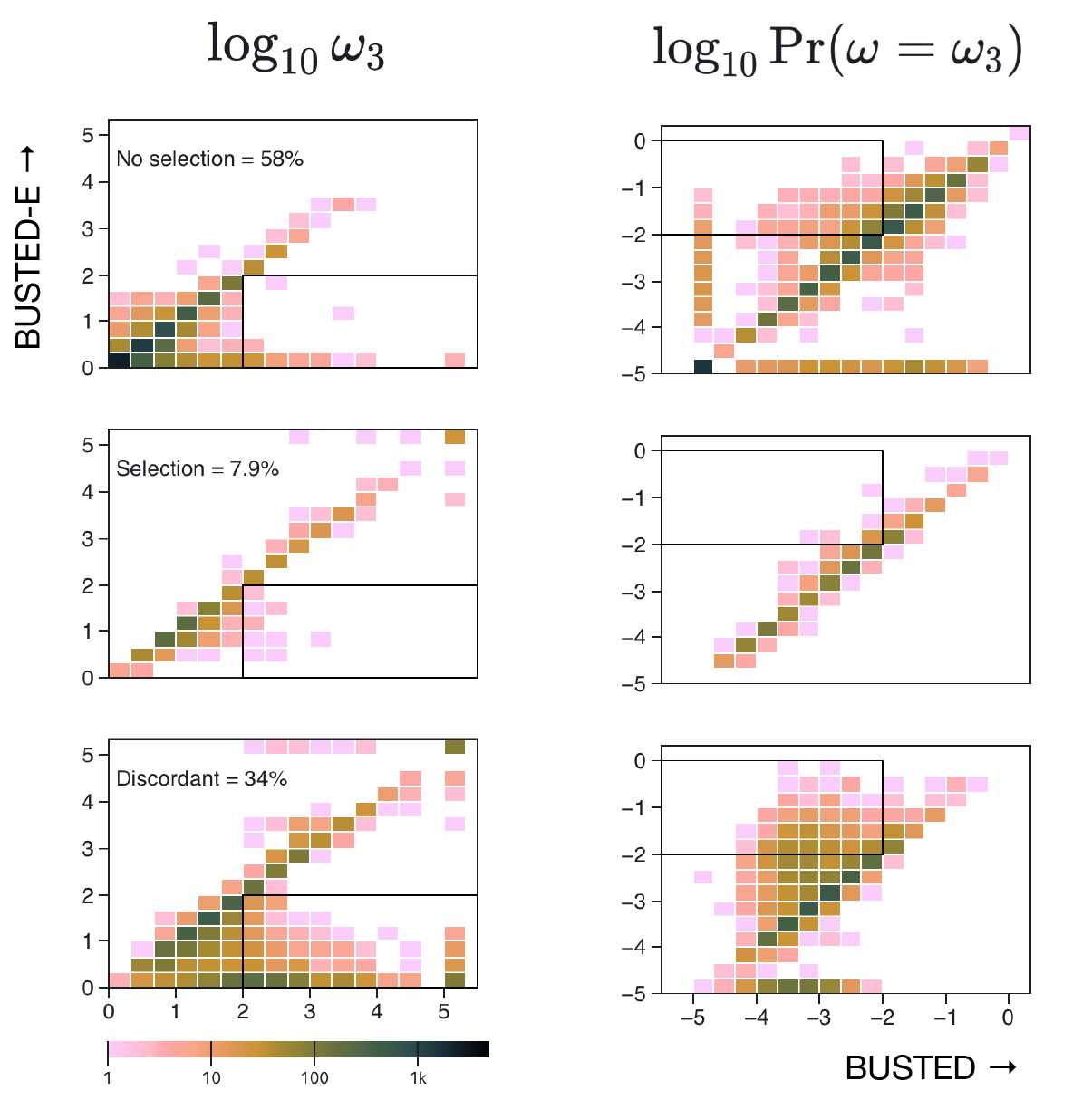
The distribution of *ω*_3_ (positive selection class estimate) estimated by BUSTED and BUSTED-E from individual genes in the Shultz and Sackton dataset ***Shultz and Sackton (2019)***. Genes are separated into groups by how the two methods classify EDS: both ‘no selection’, both ‘yes’, or ‘discordant’ top, middle and last row respectively. (Corresponding fractions of genes are shown.) The first columns shows the *ω*_3_ values the second column shows the corresponding weight of the *ω*_3_ class. For readability, values are capped at log_10_(*ω*_3_) = 5 and log_10_ Pr(*ω* = *ω*_3_) = –5 The rectangles show areas of the parameter space where BUSTED estimates are “error-like” (high *ω*, low weight) and BUSTED-E estimates lie in the more nominal range. These ranges are more densely represented among the discordant genes (last row).

### The inferred error component is small

Across all four datasets, BUSTED-E allocates an error class weight well below 1% on average, typically less than 0.1% (***Table 2***). The majority of the alignments (with one exception) have an estimated error weight of exactly 0, i.e., the BUSTED-E model effectively reduces to the simpler BUSTED model. A median of 9-32 individual codons per alignment are marked for masking by BUSTED-E (see Methods).

Few alignments have a strict statistical preference for BUSTED-E. BUSTED-E and BUSTED can be tested using an LRT which accounts for boundary conditions (e.g., when *p*_*e*_ = 0, *ω*_*e*_ is unidentifiable, ***Self and Liang*** (***1987***)). We chose to use a conservative ^2^ test statistic with two degrees of freedom (***Murrell et al., 2015***). Between 0.5% and 6.9% of the alignments reject BUSTED in favor of BUSTED-E (*p* f 0.05, ***Table 2***). For these datasets, the presence of an error component provides a significantly better fit to the data. However, on the majority of alignments where the two models disagree (BUSTED finds evidence of EDS, and BUSTED-E does not), BUSTED cannot be rejected in favor of BUSTED-E. For the Shultz and Sackton (***Shultz and Sackton, 2019***) dataset, for example, of the 3,814 alignments where BUSTED finds EDS (at *p* = 0.05), there are 3,552 (93.2%) cases when BUSTED cannot be rejected in favor of BUSTED-E (at *p* = 0.05).

### Better performance of the BUSTED-E null explains much of the discordance between the models

Unlike BUSTED, where the null model for EDS testing does not include any provision for positive selection (all *ω* f 1), the error component with *ω*_*E*_ g 100 is still available for BUSTED-E. To illustrate, in the SUGP2 (SURP And G-Patch Domain Containing 2) gene alignment from Shultz and Sackton (***Shultz and Sackton, 2019***) (Supplementary Table 2), BUSTED shows strong evidence for EDS (p *<* 0.0001), but BUSTED-E does not (*p* = 0.12). The alternative models for EDS tests have exactly the same log-likelihood scores and rate estimates (there is no error estimated component), but for the null model, BUSTED-E allocates 0.05% weight to the error component. The specific sites which are most affected by this change are those where multi-nucleotide substitutions (MNMs, e.g. CAT : AGA) are inferred to occur. Several recent studies report evidence that MNMs affect tests for EDS, increasing false positive rates when standard models are used (***Venkat et al., 2018; Dunn et al., 2019; Lucaci et al., 2023***). BUSTED-E can be extended to model MNMs directly (discussed below), although this is not our main focus here. Second, consider another discordantly classified gene, COPB1 (COPI Coat Complex Subunit Beta 1, Supplementary Table 3). Here, both BUSTED (EDS p *<* 0.0001) and BUSTED-E (EDS *p* = 0.5) estimate large *ω*_3_ values for very small fractions of otherwise highly conserved alignments, which obviates the need of a separate error component for unrestricted BUSTED-E. However, the BUSTED-E null model can simply absorb this high rate in the error class, while constraining *ω*_3_ f 1. The sites where BUSTED draws the selection signal are also enriched for MNM.

### Qualitative categorization of discordant alignments

We bin all alignments where BUSTED returns a positive EDS classification, and BUSTED-E does not into five qualitative categories:

1. Statistically supported error component, defined as a significant LRT test which rejects BUSTED in factor BUSTED-E. These are the alignments that should be the primary focus for error filtering, because they contain selection signal (non-empty positive selection *ω*_3_ component in BUSTED) and an error signal (non-empty and statistically supported error *ω*_*E*_ component in BUSTED-E), e.g., gene PXDNL from ***figure Supplement 1***.
2. Possible model misspecification, defined as having an empty error class (weight of *ω*_*E*_ is 0), a positive selection component (*ω*_3_ *<* 100), and an insignificant BUSTED v BUSTED-E LRT test. For these alignments, all the selection signal from BUSTED is absorbed by the error class in BUSTED-E null (e.g., gene SUGP2, Supplementary Table 2). Thus, the evolutionary process giving rise to substitution patterns interpreted as EDS by BUSTED generates abiological substitution rates which may instead reflect model inadequacy.
3. Only abiological selection rates, defined as having an empty error class (weight of *ω*_*E*_ is 0), a positive selection component (*ω*_3_ g 100), and an insignificant BUSTED v BUSTED-E LRT test. For these alignments there is no statistical support for anything other than an abiologically high evolutionary rate (e.g., gene COPB1, Supplementary Table 3).
4. Weak basis of selection support, defined as having a non-empty error class (weight of *ω*_*E*_ is > 0), a small (*<* 0.01) weight for the selection component (*ω*_3_ g 100) in BUSTED, and an insignificant BUSTED v BUSTED-E LRT test. These datasets are similar to class type 1 defined above, but fail to reject BUSTED vs BUSTED-E.
5. All others: alignments that do not belong to any of the previous categories. These include borderline cases (BUSTED p-value near the significant thresholds), or alignments where BUSTEDE may have suffered power loss (see the simulation section).

### Power loss suffered by BUSTED-E on null data is not sufficient to explain reduced detection rates

We compared BUSTED and BUSTED-E on 6, 000 alignments simulated under BUSTED using varying alignment sizes, divergence levels, and *ω* distributions. We found that BUSTED-E suffers some power loss when the underlying data are generated using a model without the error component. The loss is minimal for cases where the selection is strong. BUSTED-E shows a more significant power loss when selection is weaker, but that also occurs with BUSTED. Model averaging recovers much of the power lost due to the unnecessary error component for all scenarios (see ***Figure 5***). BUSTED-E does not introduce additional systematic biases to parameter estimates (see ***figure Supplement 1***). Thus, we conclude that the dramatic reduction in EDS detection rate cannot be primarily attributed to power loss.

**Figure 5.**
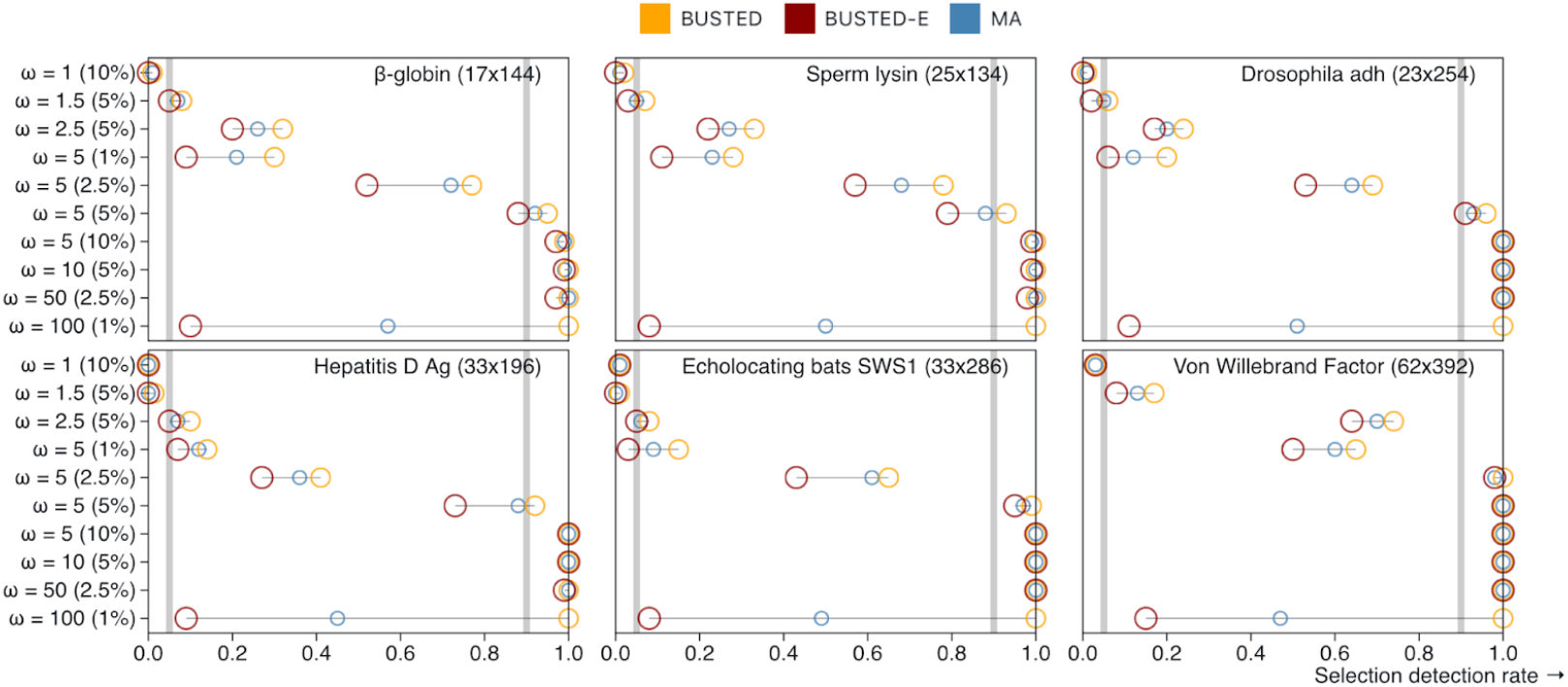
Rates of EDS selection on data simulated with the BUSTED model (Supplementary Table 6). A LRT test with pf0.05 constitutes a positive result. Horizontal reference lines demarcate 0.05 and 0.90 rates. The datasets are sorted by the number of characters, smallest to largest. Circles are of different sizes to eliminate overlap, and sizes have no specific meaning. **Figure 5—figure supplement 1**. Rates of EDS selection on data simulated with the BUSTED model (Supplementary Table 6). A LRT test with pf0.05 constitutes a positive result. Horizontal reference lines demarcate 0.05 and 0.90 rates. The datasets are sorted by the number of characters, smallest to largest. Circles are of different sizes to eliminate overlap.

### Characterizing BUSTED-E alignment filtering

BUSTED-E can annotate individual codons in the alignment with a score corresponding to the confidence that it is attributable to the error class. Codons with high error-class scores can be masked with gaps and filtered alignments then passed on to other tools.

When applied to the empirical alignment collections, BUSTED-E filtering masks a modest fraction of input alignments (***Table 3***). For the substantial majority of alignments, the filtering procedure does nothing. For the alignments where at least one codon is filtered, an average of 2-3 codons per 1000 codons are masked. With the exception of Wu et al. ***Wu et al***. (***2018***) (gapless input alignments), the filtering/masking procedure adds one new masked codon per 10-20 existing gapped codon positions.

**Table 3.** BUSTED-E filtering procedure statistics. Unaffected alignments are those where no filtering is deemed necessary. All the following statistics are computed only over those alignments where some filtering was done. Masked codons per is the median [2.5% - 97.5%] number of codons masked per: alignment (raw counts), a single column (normalized), 1000 codons of sequence (normalized). As fraction of total gaps, refers to the median [2.5% - 97.5%] percentage of the gaps in the filtered alignment that are due to those added by filtering (the others were already there). *Input datasets for this reference contained no gaps.

| Study | Unaffected alignments (%) | Masked codons per |  |  | As fraction of total gaps (%) |
| --- | --- | --- | --- | --- | --- |
|  |  | Alignment | Site | 1k of sequence |  |
| Schneider | 6263 (66.6) | 11 [1.0-79.1] | 0.0027 [0.00022-0.011] | 2.7 [0.22-11] | 7.7 [0.42-96] |
| Nguyen | 3172 (63.7) | 9 [1.0-71.0] | 0.0028 [0.00026-0.014] | 2.8 [0.26-14] | 5.3 [0.31-100] |
| Wu | 3771 (88.8) | 9 [1.0-52.7] | 0.0023 [0.00013-0.011] | 2.3 [0.13-11] | 100 [100-100]* |
| Shultz | 7245 (64.3) | 32 [1.0-262.0] | 0.0019 [7.5e-05-0.0095] | 1.9 [0.075-9.5] | 9.3 [0.24-100] |

Next, consider the effect that filtering has on EDS detection with standard BUSTED (no error component) models (***Table 2***). For all 4 datasets, when BUSTED is applied to filtered datasets, EDS detection rate is reduced significantly but remains higher than BUSTED-E on original datasets in 3 of 4 cases. In these 3 cases, BUSTED-E and MA approaches have essentially the same detection rate on original/filtered data — an indication of internal consistency. The dissenting dataset has an unusually high fraction of original alignments where BUSTED-E is preferred to BUSTED by LRT (0.2), and these alignments are gappy (5% of total alignment size comprises gaps, see ***Table 1***). Indeed, masking the codons most significantly contributing to the error signal in the original data reduces the residual *ω*_*E*_ weight (last column of ***Table 2***) several fold, with *>* 90% of the datasets allocating no weight to this model component (e.g., ***Figure 1***).

### BUSTED-E filtering compared to existing methods

We applied two widely used alignment filtering methods: BMGE (***Criscuolo and Gribaldo, 2010***) and HMMCleaner (***Franco et al., 2019***) to alignments from Schulz and Sackton (***Shultz and Sackton, 2019***), and compared how they performed relative to BUSTED-E. The primary goal was to determine whether BUSTED-E targets the same error modalities as the existing methods, both of which are motivated by finding regions of sequences that have low homology to other sequences. The results of this comparison are summarized in ***Table 4***.

**Table 4.**
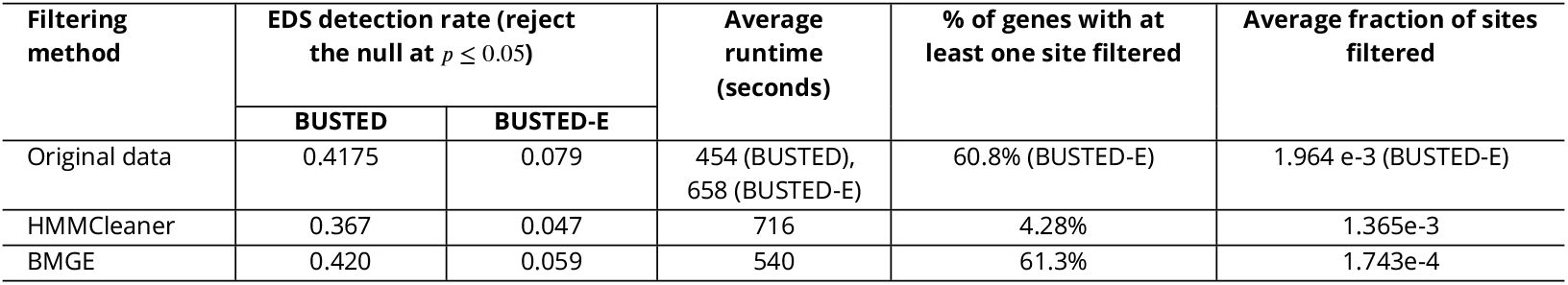
The effects of various data filtering approaches on BUSTED and BUSTED-E performance; the “original data” row also includes filtering metrics using BUSTED-E.

**Table 5.** Key BUSTED-E rate variation parameters.

| Parameter | Role | Variation | Distribution |
| --- | --- | --- | --- |
| $\alpha(s)$ | Relative synonymous rate at a site | Independent among sites. | M-bin unit mean general discrete distribution (GDD) |
| $\omega(s, b)$ | Non-synonymous to synonymous substitution rates ratio on a branch at a site | Independent among sites and branches. | N-bin GDD. $\omega_1 \leq \omega_2 \dots \leq 1 \leq \omega_{N-1} \ll E \leq \omega_N$ |

HMMCleaner and BMGE target different errors than BUSTED-E. This follows from (a) noticing that BUSTED found EDS at about the same rate (∼ 30 – 40%) in the alignments filtered by these methods as in the original alignments, and (b) because EDS detection with BUSTED-E was dramatically lower than BUSTED for datasets passed through HMMCleaner and BMGE. In other words, from the standpoint of BUSTED-E, most of the errors it detects pass unchallenged by the other two tools. All three methods mask very small fractions of alignments (0.02 – 0.2%). As a practical advantage, the average run times for BUSTED-E, which performs selection detection and error-filtering jointly, are comparable to the average run times of HMMCleaner and BMGE, which only perform filtering. It is important to reiterate that Schulz and Sackton (***Shultz and Sackton, 2019***) alignments had already undergone alignment filtering with custom scripts (***Jarvis et al., 2014***). Consequently, it is reasonable to assume that most of the homology, annotation, or assembly errors which are the primary targets of BMGE and HMMCleaner had already been removed and what remains are smaller errors that affect individual sites.

### BUSTED-E model extensions

The core idea of BUSTED-E is to identify alignment regions which appear aberrant in relation to the underlying evolutionary model. When this model is changed, the definition of aberrant patterns is also likely to change. Consider, for example, what happens if the codon substitution model is extended to permit instantaneous multi-nucleotide (MH) substitutions (***Lucaci et al., 2021***). As we and others have shown previously (***Lucaci et al., 2023***), including MH support in the context of selection detection dramatically lowers the rate of detection. For example, on data from Schulz and Sackton (***Shultz and Sackton, 2019***), MH-enabled models detected EDS at a rate about 3 times lower than models without MH support (***Lucaci et al., 2021***). Much of this reduction is attributable to the difference in how multi-nucleotide substitutions that occur along short branches are handled. For standard models, MH substitutions drive up the estimates of *ω*, and in many cases one or a few of these substitutions is all that lends statistical support to positive selection in an alignment. For MH-enabled models these types of substitutions will be accommodated by separate 2- or 3-hit substitution rates, and reduce *ω* estimates. As an illustration, we include examples of datasets which are discordant under BUSTED and BUSTED-E (Supplementary Table 4), but not discordant under +MH versions of this model from each of the five qualitative categories we defined previously. Thus, multi-hit and error model components are interacting, with the change in the baseline model substantively modifying what data features are considered “error-like”.

We reran the alignments from Schulz and Sackton (***Shultz and Sackton, 2019***) with the BUSTED- MH model as the baseline, adding the error-sink component, and made the following observations.

1. The inclusion of +MH reduces EDS detection rates about four-fold, to 10.1% with BUSTED and to only 1.8% with BUSTED-E.
2. The datasets where BUSTED (without MH) finds EDS and BUSTED-E (also without MH) does not, are somewhat enriched for datasets where there is statistical evidence for including MH support (Odds Ratio = 1.2, p *<* 0.001). The addition of MH support to BUSTED abrogates EDS detection rates. For example, 20% of the datasets assigned to Type 2 discordant class (“Possible model misspecification”) are also discordant between BUSTED and BUSTED-MH, with discordant datasets showing elevated rates of multi-nucleotide substitutions compared to the negative datasets (***Figure 6***). Interestingly, concordant positive datasets also have MH rates elevated compared to concordant negative datasets (***Figure 6***).
3. There is an interaction between MH rates and the presence of the error component; when a non-zero error component is inferred, 2- and 3-nucleotide rate estimates are lowered (Supplementary Table 5).
4. There is a significant additional computational cost for adding the -MH component. On average, BUSTED-MH run times are on average 8x longer than BUSTED run times, and BUSTED-MH-E run times are on average 9x longer than BUSTED-E run times.

**Figure 6.**
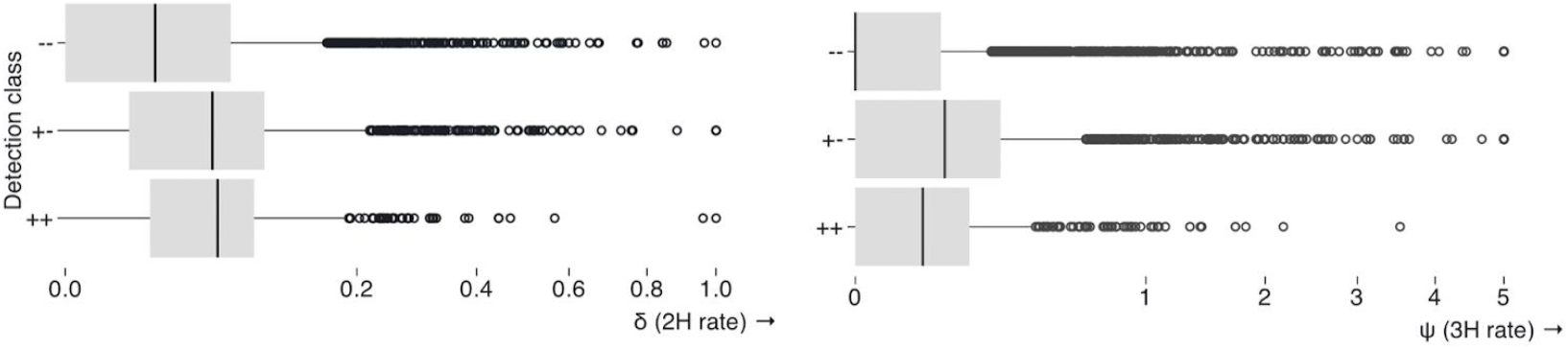
Point estimates of multiple-nucleotide substitution rates (under BUSTED-MH) stratified by BUSTED/BUSTED-E (without MH support) EDS classification at pf0.05, e.g. +-: detected as EDS by BUSTED, but not by BUSTED-E.

## Discussion

In the early days of comparative evolutionary analyses, finding a positively selected gene was rare and remarkable, because data were sparse and statistical methods were insensitive, requiring truly powerful signals (***Hughes and Nei, 1989; Lamers et al., 1993***). Today, there are thousands of sequenced genomes, and statistical methods are much more sensitive. As a result, analyses often find that the majority of genes may have been subject to EDS (***Shultz and Sackton, 2019***). The imprimatur of “positive selection” has lost its luster. Researchers must further refine prolific candidate lists of selected genes to confirm that the findings are robust and meaningful. Multiple methods are applied, and only genes where all methods agree are tagged as selected. Complex data filtering procedures are devised and run to reduce the influence of outliers and poor-quality sequences.

Many alignment filtering and curation tools and protocols exist, including GBLOCKS (***Castresana, 2000***), ALISCORE (***Misof and Misof, 2009***), TrimAl (***Capella-Gutiérrez et al., 2009***), BMGE (***Criscuolo and Gribaldo, 2010***), Zorro (***Wu et al., 2012***), GUIDANCE (***Sela et al., 2015***), HMMclean (***Franco et al., 2019***), HoT (***Landan and Graur, 2007***). However, since the the correct homology and alignment is not known in practice, all approaches must rely on reasonable, but ultimately unverifiable assumptions about what constitutes error. These assumptions are usually not integrated with or informed by downstream analyses that consume MSAs. This creates the potential for errors that bias downstream analyses to leak through the filters. As we demonstrated using published data for which the authors made diligent effort to filter alignment errors with modern tools, “obvious to the eye” (***Figure 1***) errors still made it through the filtering process. Sensitive branch-site dN/dS models like BUSTED have little trouble pinpointing such errors with aberrant parameter estimates: high dN/dS values affecting tiny fractions of the alignment. Leveraging this simple intuition to define *ad hoc* error-regions for dN/dS parameter estimates (dN/dS g 100, weight f 1%) turns out to be quite effective at reducing the rate of positive selection detection, with the implication that most of the detected EDS genes in a typical large-scale screens are false positives.

For a typical practitioner, dN/dS tests of selection are “black boxes”, and the source of signal, or the influence of outliers is difficult to assess ***Spiess et al. (2024)***, especially if applied at scale. Intuitively, inference of selection is more robust if it is based on multiple substitutions across multiple sites. As sequencing projects ramp up species coverage, including many divergent species in selection analysis increases the chance of misalignment and other error. Given enough data with some errors, and the challenges of catching such errors via automated procedures, every gene in the genome would be inferred under positive selection on at least one branch eventually. This is biologically meaningless and makes selection analyses useless as a binary discriminative tool. On the other hand, even for moderate-sized datasets analyzed here, the BUSTED-E error correction framework improves downstream biological inference by reducing false positive signals due to confounders such as protein length and improving functional relevance (***Figure 3***).

We suggest that whenever researchers consider using dN/dS analyses, especially in high-throughput settings, an error-correcting framework like BUSTED-E be used routinely. In genome-wide scans, especially those involving a large number of species, an automated method to account for sequencing and alignment error will be crucial, lest all of the signal be swamped with residual error. BUSTED-E is not meant to be a primary filter for gross or large-scale errors, which are better handled by other tools, but rather as a secondary filter, as done here. To guard against the loss of power, we recommend using model-averaged approaches. While alignment masking is not necessary for BUSTED-E to mitigate errors during selection screens, its ability to perform such masking is useful prior to applying other tools not equipped with error filtering (***Kosakovsky Pond and Frost, 2005; Yang, 2007; Murrell et al., 2012***).

BUSTED-E is at its heart an *ad hoc* method. On the one hand, it finds many obvious errors missed by routine filtering, reduces EDS detection rate to credible levels, and results in more sensible functional description of selected genes. On the other hand, the error component is heuristic and phenomenological, albeit tunable. Other settings for the error component class will influence what is considered an error. Similarly, the choice of baseline substitution model has a major impact on error filtering. We view BUSTED-E as a practical and necessary solution to deal with pervasive alignment contamination to be refined and improved. Because we do not yet fully understand the distribution and nature of sequencing and alignment errors, we expect that BUSTED-E will fail to detect some real errors and also erroneously flag some biological variation as error. Power loss, possibly significant for some selection regimes, is also a concern. Encouragingly, our simulations show that given even moderate selection signal, power loss is minimal and can be mitigated with model averaging. Despite these limitations, BUSTED-E is a robust and scalable “drop-in” solution for improving the accuracy of evolutionary selection analyses in the presence of alignment errors, contributing to a more nuanced understanding of natural selection and adaptive evolution.

## Methods

### Model description

BUSTED-E (Branch-site Unrestricted Test for Episodic Diversification with an Error component), is a random-effects model of codon evolution, which builds upon the BUSTED class (***Murrell et al., 2015; Wisotsky et al., 2020; Lucaci et al., 2023***). BUSTED-E is designed to test for a fraction of the alignment evolving with dN/dS (*ω*) *>* 1, i.e. subject to positive diversifying selection, while also accommodating an “abiological”, or error-like, mode of evolution. The key parameters describing the evolution at alignment site *s*, along the branch of a phylogenetic tree *b* are the instantaneous rate of synonymous substitutions *a* and the ratio of non-synonymous to synonymous substitutions *ω*, respectively.

The key feature of BUSTED-E is the inclusion on an error-component to the distribution of *ω*, with its ratio value bounded below by a large (*a priori* specified) number E, e.g. *E* = 100, and the maximal weight that can be allocated to this error component bounded from above by *p*_*e*_, e.g. 1%. Exceedingly large *ω* values are a hallmark of unrealistic deviations from the expected evolutionary process, and as such are categorized as alignment or sequencing error. We chose these specific values because they represented specific modes of error in empirical data (***Table 2***), and because they were decimal round numbers.

Error class parameters can be adjusted, and potentially even inferred from the data as model parameters. BUSTED-E is otherwise identical to the previously described and validated BUSTED-S model. The default, user-tunable, hyperparameter values are *N* = 4 (number of *ω* bins) and *M* = 3 (number of *a* bins). For N=4, the *ω* distribution accommodates two classes for negative selection or neutral evolution (*ω*_1_ f *ω*_2_ *<* 1), a class for positive selection or neutral evolution (1 f *ω*_3_), anda class for errors (100 = *E* f *ω*_4_).

### Formal model definition

The BUSTED-E model is a standard continuous time discrete space Markov model of codon substitution, which partitions codon substitutions into three classes, with the corresponding instantaneous rate matrix 𝒬, parameterized following the Muse-Gaut (MG94) (***Muse and Gaut, 1994***) approach. Here, *θ*_*ij*_ = *θ*_*ji*_ denote five estimated nucleotide-level substitution bias parameters (*θ*_*Ag*_ = 1 for identifiability), 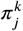 are the corrected estimates of position-specific equilibrium frequencies for the target nucleotide (***Pond et al., 2010***). As described above, *a, ω* vary from site-to-site and from (branch,site) to (branch,site), respectively. All parameters, including branch lengths, are estimated using maximum likelihood, as implemented in the HyPhy package v2.5.56 or later (***Kosakovsky Pond et al., 2020***).

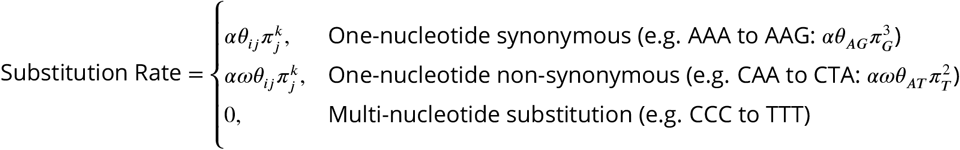

### Statistical testing using BUSTED-E

BUSTED-E can be constrained down to BUSTED-S, either as *ω*_3_ = *ω*_*E*_ or its *p*_*e*_ = 0. Therefore, we use the conservative χ^2^ distribution with two degrees of freedom to compute LRT p-values for rejecting BUSTED-S in favor of BUSTED-E (***Self and Liang, 1987***). We test the alignment for evidence of EDS by constraining *ω*_*N*–1_ = 1, optimizing this null model, and comparing it to the alternative BUSTED-E (where *ω*_*N*–1_ g 1) using the likelihood ratio test. Once again, we use the conservative χ^2^ with two degrees of freedom as the asymptotic distribution of the test statistic to assess significance. Because the addition of an error rate class is expected to reduce statistical power when the correct model has no error component, or when the alignment is “error-free”, we compute a model-averaged test of significance (following the general framework described in (***Lucaci et al., 2023***)). We fit both BUSTED-S and BUSTED-E to the same data, compute the Akaike weights for each model using the small sample AIC_c_ information criterion score, and weigh LRT p-values for EDS by the corresponding Akaike weights: *p*(*MA*) = *w*(BUSTED[S])*p*(BUSTED[S]) + *w*(BUSTED-E)*p*(BUSTED-E). An Akaike weight of the model is defined as exp 0.5(min *AIC*_c_ – model*AIC*_c_), with the minimum taken over the models being compared, and normalized to sum to 1 over all models.

Model averaging is a way to control potential power loss. Assuming the worst-case scenario, where BUSTED-E offers no improvement in log-likelihood at the cost of two additional parameters, its AIC_c_ score will be 4 or more units higher than the AIC_c_ score for BUSTED-S. Further assume that *p*(BUSTED-E) = 0.5 (the highest p-value for one-sided tests as defined in BUSTED). The contribution from BUSTED-E to the model-averaged p-value will be 0.5 × *r*(BUSTED-E) f 0.06.

### Model variations

The BUSTED-E concept can be extended in several directions. First, we could alter the baseline model, for example by removing the synonymous rate variation (SRV) component, or adding a component for accommodating multi-nucleotide substitutions (***Lucaci et al., 2023***). Because the underlying model specifies what the expected biological reality is, it will influence the inference of what constitutes possible alignment errors. Second, we could partition the tree into “foreground” and “background” branches, as is commonly done for detecting lineage-specific selection, and apply to them different *ω* distributions and error components; these options are already implemented and available within HyPhy, but not discussed in detail further.

### Heuristic error filtering

We implement the following heuristic for alignment filtering. Our procedure applies to codons at individual branches, and consists of “masking” a subset of codons with gap characters (–-). First, for each branch and site in the alignment, we compute two empirical Bayes factors (EBF) asking (1) if the *ω* value comes from the error class (EBF_e_), and (2) if the *ω* value comes from the (non-error) positive selection class (EBF_s_). To obtain those EBFs, we compute, for each (branch, site) pair *N* conditional probabilities 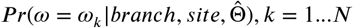, with 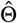 used to represent all other ML parameter estimates. This involves the calculating *N* site-level phylogenetic likelihood functions, and setting the weight of *ω*_*k*_ to 1 on a particular branch. These conditional probabilities can be converted into empirical Bayes posterior probabilities, simply by dividing each by 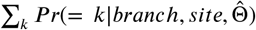, and computing EBF as the ratio of posterior to prior odds of *ω* = *ω*_*k*_. The computational cost of the procedure for all (branch, site) pairs is roughly equivalent to *N* full-alignment likelihood calculations, and is negligible compared to the cost of BUSTED-E model fitting. EBF_e_ is computed for the events *ω* = *ω*_*N*_ (error class) vs *ω ≠ ω*_*k*_ (not error class). EBF_s_ is computed for the events *ω* = *ω*_*N*_ (error class) vs *ω* = *ω*_*N*–1_ (selection class).

A (branch, site) will be filtered when the following two conditions are satisfied. 1. EBF_e_ g *C*_1_, where *C*_1_ *>* 1, default = 100 is a cutoff set *a priori* by the user, with higher values being more stringent (more conservative filtering). This condition encodes sufficient evidence that (branch,site) belongs to the error class. 2. EBFs g *C*_2_, where *C*_2_ *>* 1, default = 20 is a cutoff set *a priori* by the user, with higher values being more stringent (more conservative filtering). This condition encodes sufficient evidence that the error class is strongly preferred to the positive selection class.

When the branch being filtered is a leaf, only the corresponding (observed) codon is masked. For an internal branch, there is no corresponding observed codon, and all of the observed codons in the smaller of the two tree splits defined by the branch are masked. A third user controlled threshold 0 *< C*_3_ f 1, default = 0.4 is used to mask the entire site (as potentially unreliable), if more than *C*_3_ fraction of observed codons have been masked. ***Figure 7*** illustrates the filtering procedure on a few key examples.

**Figure 7.**
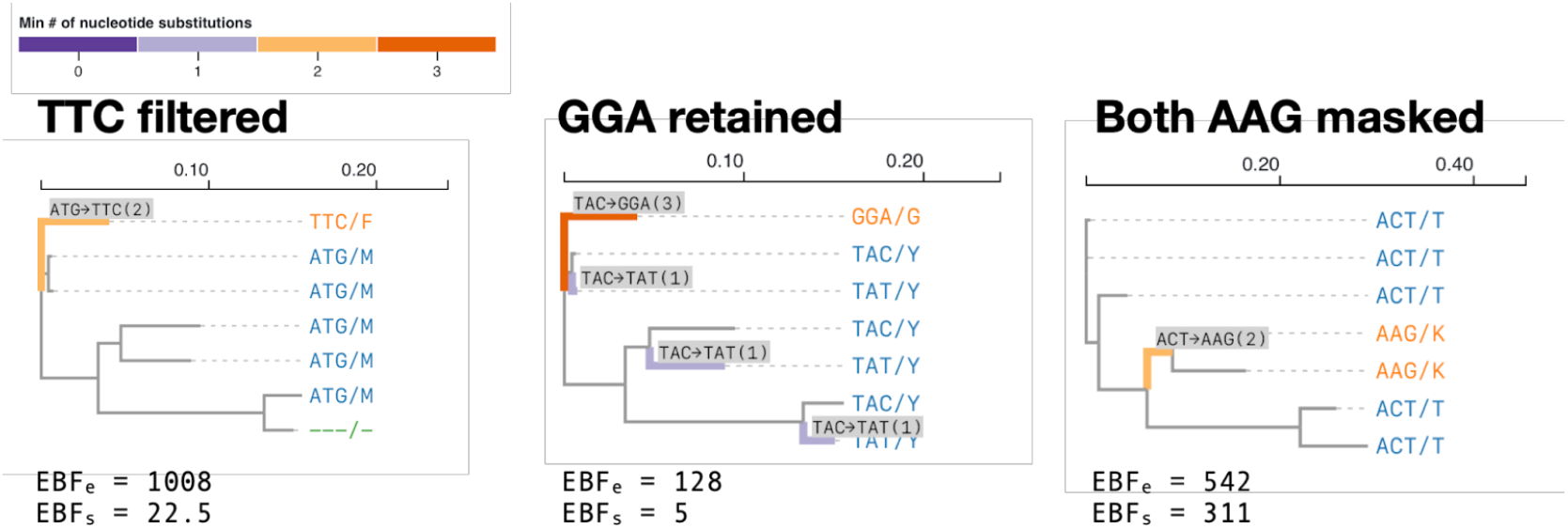
Examples of codon sites filtered and retained by the heuristic filtering procedures based on the empirical Bayes factors for error class membership and for positive selection.

### Implementation and performance

BUSTED-E is implemented in HyPhy versions 2.5.53 or later, as well as on the Datamonkey web application ***Weaver et al. (2018)***; versions of BUSTED-E are different between HyPhy versions, and all of the analyses in the paper were done with 2.5.60, 2.5.61, or 2.5.62. Supplying –error-sink Yes to a hyphy busted call turns on the error sink component. Because BUSTED-E is a mixture random effects model, its parameter estimation using direct optimization (as done here) is sensitive to starting conditions. We devised a multistage model fitting procedure which attempts to identify good starting points to speed up BUSTED-E convergence and improve run-to-run reproducibility:

1. Fit the nucleotide GTR model to obtain initial estimates of branch lengths and nucleotide bias parameters (*θ*_*ij*_).
2. Fix all estimates from phase 1, and estimate mean *ω*_0_ value (or values if there is more than one branch partition) using the MG94xREV codon model.
3. Use parameter estimates from phase 2 as the starting point, to re-estimate branch lengths, nucleotide bias parameters, and the mean *ω*_0_ value(s) under the full MG94xREV codon model.
4. Fix parameter estimates from phase 3. Generate K (=250 by default) Latin Hypercube random samples of *ω* distributions with means constrained to *ω*_0_ estimates from phase 3, and *a* distributions with means constrained to 1.
5. Compute the phylogenetic likelihood function on the K initial points, select P (=5 by default) best scores.
6. Perform “quick-and-dirty” *ω* and *a* distribution optimizations from using the P starting points from phase 5. The optimizations are done using the Nedler-Mead simplex algorithm ***Nelder and Mead (1965)***, with the stopping criterion that consecutive iterations fail to improve the log likelihood score by at least *max*(0.5, –10^−4^ log *L*(*phase*3)).
7. Select the estimates yielding the best likelihood score as the initial condition for the full direct optimization of the BUSTED model.

Additionally, if both BUSTED and BUSTED-E are fitted to the same alignment, we first run BUSTED, and add the parameter estimates obtained from BUSTED as 50% of the starting points candidate for step 4 in the fit for BUSTED-E, with the values of the error-sink drawn class drawn randomly.

### Simulated null data analysis

We parametrically generated simulated alignments under the BUSTED (+S) model, using six empirical alignments as guides. Specifically, we fitted the BUSTED (+S) model with 2 or 3 rate classes to the biological alignments previously analyzed for EDS in multiple papers. The alignments included different numbers of sequences, varied in length, divergence, nucleotide composition, nucleotide substitution biases, and the inferred site-to-site distribution of synonymous rates (Supplementary Table 6). For each empirical dataset, we generated 100 replicates under one of 10 selective profiles (Supplementary Table 7), yielding 6,000 total alignments. We then ran BUSTED and BUSTED-E on each replicate and tabulated the results, focusing on the power to detect selection, biases in rate estimates, and the rate at which BUSTED is rejected in favor of BUSTED-E.

### Interaction with other filtering methods

To understand the type of alignment errors detected by BUSTED-E filtering, we explored how traditional alignment filtering methods interact with BUSTED-E. We selected two highly cited “conventional” alignment filtering methods: BMGE (***Criscuolo and Gribaldo, 2010***) and HMMCleaner (***Franco et al., 2019***) to use in conjunction with BUSTED-E. BMGE utilizes a sliding window approach to detect regions of high entropy, while HMMCleaner generates an alternative sequence aligned with the original sequence to find regions of low similarity. Using data from Schulz and Sackton (***Shultz and Sackton, 2019***), we filtered the alignments with each of the conventional filtering methods and used both BUSTED and BUSTED-E to explore the differences in EDS detection rate.

## Acknowledgments

This work was supported in part by the following awards: NIH/NHGRI (HG009299), NIH/NIGMS (GM151683), and NIH/NIAID (AI183870).

