## Supplementary figures and tables for "Minus the Error: Testing for Positive Selection in the Presence of Residual Alignment Errors"

### Supplementary Material

| **Study** | **Alignment** | | | **Tree length, subs/site** | **Gaps per** | | **Alignment** | **MSA filtering** |
|---|---|---|---|---|---|---|---|---|
| | **Count** | **Seqs.** | **Codons** | | **Column** | **1k codons** | | |
| Schneider | 9404 | 7 [7:7] | 469 [120:1850] | 0.41 [0.17:0.86] | 0.18 [0.0:1.0] | 25.3 [0.0:141.9] | P2C, `Darwin` | Manual |
| Nguyen | 4981 | 6 [5:6] | 511 [154:1898] | 0.21 [0.09:0.50] | 0.34 [0.0:1.17] | 58.1 [0.0:203.0] | `OrthoMam v. 6` | `Gblocks` |
| Wu | 4248 | 15 [15:15] | 246 [81:832] | 0.15 [0.08:0.44] | 0.00 [0.00:0.00] | 0.00 [0.00:0.00] | `Codons/ PRANK` | `Guidance`, manual (sliding window). Manual checking for MNM |
| Shultz | 11267 | 39 [24:44] | 387 [88:1807] | 1.3 [0.5:3.9] | 0.73 [0.00:3.41] | 19.4 [0.0:97.0] | P&C, `MAFFT, PRANK` | Custom (gap fractions & patterns; sliding window) |

**Table Supplementary Table 1. Empirical data EDS analyses**. For all per-alignment quantities (sequences, codons, tree length, gaps per) we report the median value and the [2.5% : 97.5%] range. **Gaps per** sequence are normalized per 1000 codons. **Tree length** is the cumulative branch length (expected substitutions/nucleotide) under the simple (MG94xREV) codon model. Abbreviations for the **Alignment** column are as follows. P2C: translated protein sequences are aligned, then mapped back to codon sequences; P&C: protein based homology and filtering, codon-level alignment; MNM : multi-nucleotide mutations.

| **Model** | **Log (L)** | $\omega_1$ **(weight)** | $\omega_2$ **(weight)** | $\omega_3$ **(weight)** | $\omega_E$ **(weight)** |
|---|---|---|---|---|---|
| BUSTED Alternative | $-27171.8$ | 0.00 (1.7%) | 0.15 (98.2%) | 28.23 (0.12%) | N/A |
| BUSTED Null | $-27185.9$ | 0.11 (13.7%) | 0.12 (81.7%) | 1.00 (4.6%) | N/A |
| BUSTED-E Alternative | $-27171.8$ | 0.00 (1.7%) | 0.15 (98.2%) | 28.23 (0.12%) | * (0.0%) |
| BUSTED-E Null | $-27173.2$ | 0.05 (8.5%) | 0.15 (89.7%) | 1.00 (1.8%) | 100 (0.05%) |

**Table Supplementary Table 2.** Log likelihoods and estimated $\omega$ rate distributions under null and alternative EDS testing hypotheses with BUSTED and BUSTED-E for the SUGP2 (10004) gene. (*) rate not identifiable, because the corresponding weight is estimated to be 0.

| **Model** | **Log (L)** | $\omega_1$ **(weight)** | $\omega_2$ **(weight)** | $\omega_3$ **(weight)** | $\omega_E$ **(weight)** |
|---|---|---|---|---|---|
| BUSTED Alternative | $-14556.5$ | 0.009 (95.9%) | 0.013 (4.1%) | 584.3 (0.02%) | N/A |
| BUSTED Null | $-14577.5$ | 0.00 (11.2%) | 0.001 (87.5%) | 1.00 (1.2%) | N/A |
| BUSTED-E Alternative | $-14556.5$ | 0.008 (95.8%) | 0.13 (4.2%) | 771.1 (0.02%) | * (0.0%) |
| BUSTED-E Null | $-14556.5$ | 0.009 (99.98%) | * (0.0%) | * (0.0%) | 868.5 (0.02%) |

**Table Supplementary Table 3.** Log likelihoods and estimated $\omega$ rate distributions under null and alternative EDS testing hypotheses with BUSTED and BUSTED-E for the COPB1 (11294) gene. (*) rate not identifiable, because the corresponding weight is estimated to be 0.

| Dataset | Discordant N (%) | Type 1 (%) | Type 2 (%) | Type 3 (%) | Type 4 (%) | Type 5 (%) |
|---|---|---|---|---|---|---|
| **BUSTED vs BUSTED-E** | | | | | | |
| 20333182 | 2577 (27.4) | 309 (12.0) | 793 (30.8) | 243 (9.4) | 429 (16.6) | 803 (31.2) |
| 25716091 | 954 (19.3) | 365 (38.3) | 149 (15.6) | 75 (7.9) | 109 (11.4) | 256 (26.8) |
| 29953708 | 339 (8.0) | 3 (0.9) | 145 (42.8) | 78 (23.0) | 59 (17.4) | 54 (15.9) |
| 30620335 | 3814 (33.9) | 262 (6.9) | 1181 (31.0) | 545 (14.3) | 923 (24.2) | 903 (23.7) |
| **BUSTED vs Model Averaged (MA)** | | | | | | |
| 20333182 | 1727 (18.4) | 308 (17.8) | 207 (12.0) | 166 (9.6) | 423 (24.5) | 623 (36.1) |
| 25716091 | 725 (14.7) | 359 (49.5) | 33 (4.6) | 42 (5.8) | 107 (14.8) | 184 (25.4) |
| 29953708 | 243 (5.7) | 2 (0.8) | 68 (28.0) | 69 (28.4) | 59 (24.3) | 45 (18.5) |
| 30620335 | 2763 (24.5) | 256 (9.3) | 486 (17.6) | 453 (16.4) | 888 (32.1) | 680 (24.6) |

**Table Supplementary Table 4.** Classification of alignments with discordant EDS results at $p \leq 0.05$. Discordant results were categorized into five types, described in the text, based on BUSTED and BUSTED-E results.

| Model | p-value | $\omega_1$ (weight) | $\omega_2$ (weight) | $\omega_3$ (weight) | $\omega_E$ (weight) | $\delta$ (2-hit rate) | $\psi$ (3-hit rate) |
|---|---|---|---|---|---|---|---|
| **TLK2 gene (Type 1 discordant)** | | | | | | | |
| BUSTED | 0.00 | 0.00 (0.4%) | 0.18 (100%) | 1534 (0.02%) | - | - | - |
| BUSTED-E | 0.10 | 0.00 (0.7%) | 0.17 (100%) | 10.79 (0.1%) | 2089 (0.01%) | - | - |
| -MH | 0.50 | 0.16 (40%) | 0.18 (60%) | 1.00 (0%) | - | 0.03 | 0.14 |
| -MH-E | 0.50 | 0.16 (40%) | 0.18 (60%) | 1.06 (0%) | 1480 (0.004%) | 0.03 | 0.14 |
| **SUGP2 (Type 2 discordant)** | | | | | | | |
| BUSTED | 0.00 | 0.00 (1.7%) | 0.15 (98.2%) | 28.23 (0.1%) | - | - | - |
| BUSTED-E | 0.12 | 0.00 (1.7%) | 0.15 (93.2) | 28.57 (0.1%) | * (0%) | - | - |
| -MH | 0.50 | 0.15 (100%) | * (0%) | * (0%) | - | 0.08 | 0.14 |
| -MH-E | 0.50 | 0.15 (100%) | * (0%) | * (0%) | * (0%) | 0.08 | 0.14 |
| **VEPH1 (Type 3 discordant)** | | | | | | | |
| BUSTED | 0.00 | 0.11 (92.2%) | 1.00 (7.8%) | 207 (0.03%) | - | - | - |
| BUSTED-E | 0.50 | 0.11 (92.1%) | 1.00 (7.9%) | 233 (0.03%) | * (0%) | - | - |
| -MH | 0.50 | 0.12 (54.3%) | 0.17 (42.4%) | 1.00 (3.3%) | - | 0.04 | 0.18 |
| -MH-E | 0.50 | 0.12 (54.4%) | 0.17 (42.3%) | 1.00 (3.3%) | * (0%) | 0.04 | 0.18 |
| **BSDC1 (Type 4 discordant)** | | | | | | | |
| BUSTED | 0.00 | 0.00 (50.3%) | 0.36 (49.7%) | >1000 (0.02%) | - | - | - |
| BUSTED-E | 0.50 | 0.00 (50.0%) | 0.36 (50.0%) | * (0%) | >1000 (0.02%) | - | - |
| -MH | 0.50 | 0.12 (85.3%) | 0.49 (14.7%) | * (0%) | - | 0.04 | 0.05 |
| -MH-E | 0.50 | 0.08 (78.3%) | 0.54 (21.6%) | * (0%) | >1000 (0.02%) | 0.00 | 0.00 |
| **GSTK1 (Type 5 discordant)** | | | | | | | |
| BUSTED | 0.01 | 0.33 (95.2%) | 1.00 (4.2%) | 13.24 (0.63%) | - | - | - |
| BUSTED-E | 0.29 | 0.33 (95.1%) | 0.53 (3.3%) | 4.81 (1.6%) | 165 (0.05%) | - | - |
| -MH | 0.50 | 0.16 (39.3%) | 0.52 (60.7%) | * (0%) | * (0%) | 0.04 | 0.15 |
| -MH-E | 0.50 | 0.16 (43.5%) | 0.55 (56.1%) | 2.9 (0.35%) | 134 (0.04%) | 0.02 | 0.11 |

**Table Supplementary Table 5.** The two example genes from Table 1 re-analyzed with BUSTED/BUSTED-E with support for multiple nucleotide substitutions (MH).

| Dataset | N | S | T | Composition, % (A, C, G, T) | Relative nucleotide substitution rates (AC, AG=1, AT, CG, CT, GT) | Mean $\omega$ | CoV synonymous rates | Rate classes: $\alpha, \omega$ |
|---|---|---|---|---|---|---|---|---|
| $\beta$ -globin | 17 | 144 | 2.24 | 20.8, 26.1, 29.2, 23.9 | 0.71, 0.36, 0.57, 1.59, 0.51 | 0.24 | 0.35 | 2,2 |
| Sperm lysin | 25 | 134 | 2.54 | 28.5, 21.3, 24.9, 25.3 | 0.85, 0.45, 0.66, 0.70, 0.26 | 0.94 | 0.85 | 3,2 |
| Drosophila adh | 23 | 254 | 1.36 | 23.3, 28.6, 25.4, 22.7 | 0.77, 0.51, 0.79, 2.29, 0.48 | 0.09 | 0.34 | 3,2 |
| Hepatitis D | 33 | 196 | 1.76 | 30.1, 22.3, 36.9, 10.8 | 0.44, 0.56, 0.29, 1.52, 0.21 | 0.42 | 0.84 | 3,3 |
| Echolocating bats SWS1 | 33 | 286 | 1.10 | 16.4, 30.5, 26.0, 27.1 | 0.19, 0.07, 0.13, 0.73, 0.14 | 0.22 | 0.30 | 2,3 |
| *vwf* | 62 | 392 | 5.02 | 20.7, 30.3, 30.9, 18.1 | 0.32, 0.21, 0.38, 1.25, 0.19 | 0.19 | 0.42 | 3,3 |

**Table Supplementary Table 6.** The six empirical alignments used as stencils for simulations under the BUSTED model. N = number of sequences; S = number of codons; T = total tree length, subs/nucleotide site; CoV = coefficient of variation.

| **Scenario** | $\omega_0$ | $Pr(\omega = \omega_+)$ |
|---|---|---|
| 1. Neutral Evolution | 1.0 | 0.10 |
| 2. Very weak EDS | 1.5 | 0.05 |
| 3. Weak EDS | 2.5 | 0.05 |
| 4. EDS, very small fraction | 5.0 | 0.01 |
| 5. EDS, small fraction | 5.0 | 0.025 |
| 6. EDS | 5.0 | 0.05 |
| 7. EDS, large fraction | 5.0 | 0.10 |
| 8. Strong EDS | 10.0 | 0.05 |
| 9. Very high $\omega$ | 50.0 | 0.025 |
| 10. Error only | 100.0 | 0.01 |

**Table Supplementary Table 7.** Parametric BUSTED simulation scenarios (no error).

---

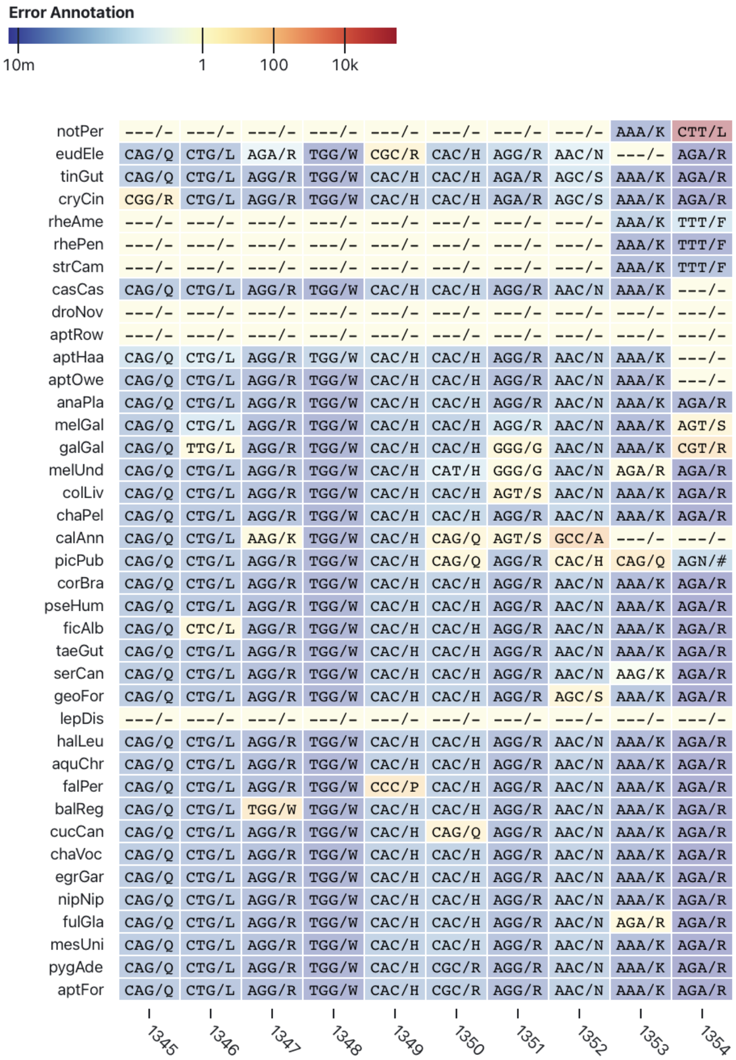


**Figure 1—figure supplement 1.** Apparent local misalignment at the 3' end of the PXDNL gene from Shultz and Sackton (***Shultz and Sackton, 2019***). The color palette indicates empirical Bayes factor support for assignment to the error class.

664

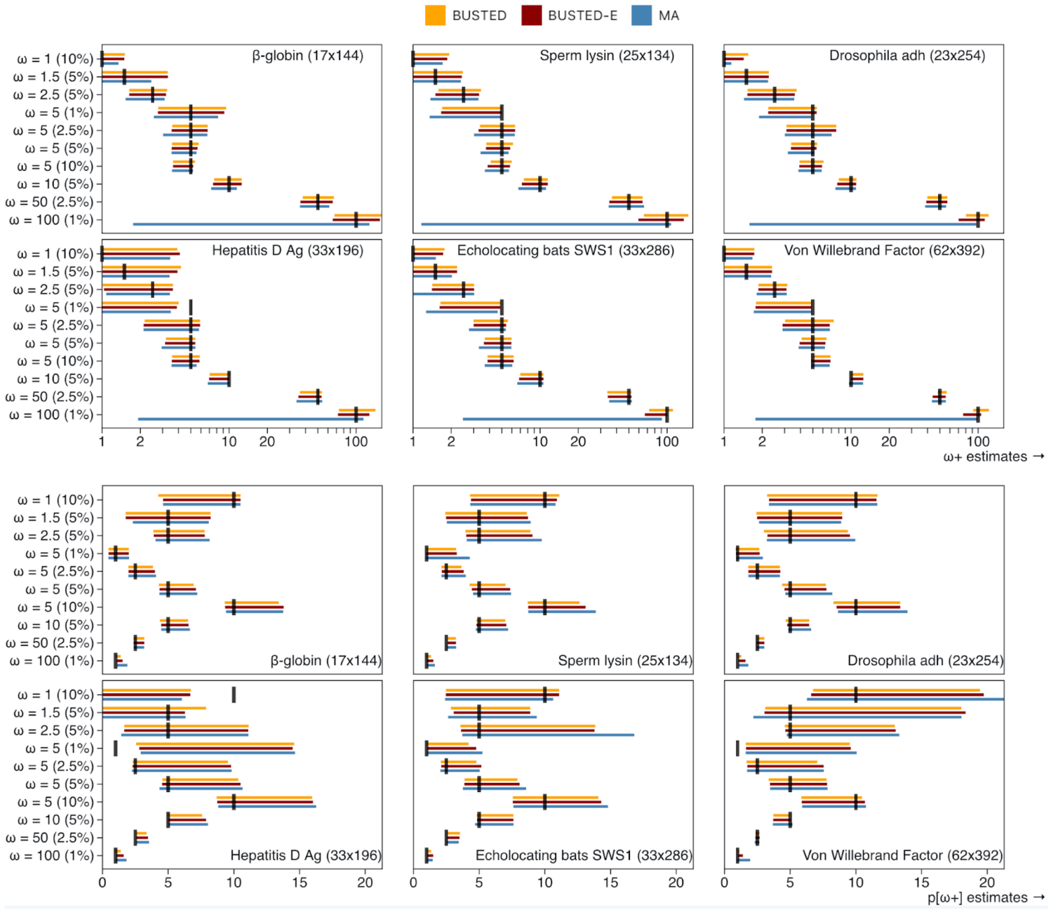


**Figure 5—figure supplement 1.** Rates of EDS selection on data simulated with the BUSTED model (Supplementary Table 6). A LRT test with p≤0.05 constitutes a positive result. Horizontal reference lines demarcate 0.05 and 0.90 rates. The datasets are sorted by the number of characters, smallest to largest. Circles are of different sizes to eliminate overlap.
